# Disruption of the developmental factor Otp in the adult forebrain reveals its diverse physiological functions

**DOI:** 10.1101/2025.09.09.675064

**Authors:** Maayan Tahor, Yael Kuperman, Tali Nahum, Michael Tsoory, Batya Bejar, Estar Regev, Janna Blechman, Jakob Biran, Alon Chen, Gil Levkowitz

**Affiliations:** Department of Molecular Neuroscience, Weizmann Institute of Science, PO Box 26, Rehovot 7610001, Israel; Department of Molecular Cell Biology, Weizmann Institute of Science, PO Box 26, Rehovot 7610001, Israel; Department of Veterinary Resources, Weizmann Institute of Science, PO Box 26, Rehovot 7610001, Israel; Department of Brain Science, Weizmann Institute of Science, PO Box 26, Rehovot 7610001, Israel; Department of Poultry and Aquaculture, Agricultural Research Organization, Rishon Lezion, 7528809, Israel

## Abstract

Orthopedia (Otp) transcription factor is a critical determinant in the development of the neuroendocrine hypothalamus, and its embryonic deletion results in lethality. Although Otp expression is maintained throughout life, its physiological function in adulthood is not well understood. Here, we generated a forebrain-specific, tamoxifen-inducible, conditional knockout mouse model to investigate the roles of Otp beyond development. Conditional deletion of Otp in two-month-old mice resulted in impaired stress responses, characterized by increased depressive-like behavior and elevated stress-induced cortisol levels. It also led to various metabolic changes, including reduced thyroid hormone levels and body temperature, a higher percentage of fat mass, and diminished responsiveness to ghrelin without affecting food intake, energy expenditure, or body weight. This composite metabolic phenotype was associated with reduced hypothalamic neuropeptides TRH, CRH, AgRP, and NPY expression. Our findings highlight the role of Otp in adult physiological functions as a key neuroendocrine integrator of adaptive stress response and energy balance.

## Introduction

The hypothalamus integrates signals from the external and internal environments to maintain physiological homeostasis, the dynamic set-point to which the body is tuned. Multiple neuronal types, localized to distinct hypothalamic regions, are vital for properly controlling thermoregulation, defense, reproduction, and food and liquid intake (1). These intricate functions are enabled by a precise developmental program, leading to a specific anatomical microstructure and neural connectivity. Several transcription factors are important for the proper development of this microstructure. Their combinatorial co-expression defines specific neuronal subpopulations within hypothalamic subnuclei (2). Embryos missing Single-minded homolog 1 (SIM1) and Aryl hydrocarbon receptor nuclear translocator 2 (ARNT2) lack a recognizable paraventricular nucleus (PVN), supraoptic nucleus (SON), and the anterior periventricular nucleus (aPV) (3), which eventually affects the organism’s survival. Orthopedia (Otp) is a prominent evolutionarily conserved hypothalamic transcription factor across species from invertebrates to mammals (4),(5). Otp is expressed in highly conserved hypothalamic domains, including practically all neuropeptidergic neurons of the neuroendocrine hypothalamus domains (6–8). Numerous studies, including our own, have demonstrated that Otp plays a crucial role in the development of hypothalamic neurons in both mouse (9,10) and zebrafish (11–13)

Otp expression is maintained in the adult human, mouse, and zebrafish forebrain, implying a post-developmental physiological role (14). However, homozygous Otp null mice display progressive impairment of crucial neuroendocrine developmental events and consequently die soon after birth, hence impeding functional evaluation of its role in the adult.

Zebrafish express two paralogous genes, namely *otpa* and *otpb*, whose expression patterns largely overlap, and adult fish with a single mutation in either gene are viable and fertile. Consequently, we previously took advantage of this fish model and showed that adult *otpa* mutants display both anxiety-like and social deficits (15,16). However, we could not determine whether these deficits result from Otp action during development or if they reflect a genuine physiological role of Otp in the adult brain.

Several mouse studies have addressed the molecular and physiological consequences of altered *otp* expression in adulthood by studying heterozygous animals, mice bearing a mutated *otp* gene, or by site-specific deletion using the cre-lox system. Mice heterozygous for a missense mutation (pR108W), which is localized to the highly conserved homeodomain of Otp, display reduced oxytocin (OXT) expression but only a transient reduction in AgRP, thyrotropin-releasing hormone (TRH), and somatostatin (SST) expression(17). These mice are hyperphagic, obese, and have behavioral deficits (17). Similarly, the knock-in mice, carrying the Q153R/+ Otp allele, identified in an obese child, were hyperphagic and exhibited obesity that increased under a high-fat diet (18). Site-specific deletions, including the deletion of Otp in AgRP neurons using an inducible Agrp-cre, led to fewer AgRP and (15) SST-expressing neurons without obvious gross abnormalities (19). Additionally, mice lacking one copy of Otp in Sim1 neurons gradually gain more weight when consuming chow, and this effect is enhanced on a high-fat diet (18). Similarly, a few cases of humans with the Otp polymorphism are characterized by early-onset obesity (17,18). A recent study demonstrated that the deletion of Otp in the PVN of adult mice led to reduced SST and OXT expression, hyperphagia, and obesity (18).

Together, the data has so far indicated that a lifelong disruption in Otp expression affects the regulation of energy balance and the responses to stress in adulthood. Nevertheless, since the deletion of Otp in those studies occurred during the embryonic stage, it is difficult to clearly distinguish between the developmental effects of Otp and its functions during adulthood.

To that end, the current study has explored the roles of Otp in adult physiology by inducing deletion of its expression in the adult forebrain. The findings indicate that Otp is essential for adequate physiological and behavioral adaptations to stress in the adult brain and for the neuroendocrine regulation of various, sometimes opposing, metabolic functions.

## Results

### Generation and validation of Otp conditional knockout mice

As the developmental loss-of-function of Otp leads to near-birth lethality, germline O*tp*-KO mice are unsuitable for studying Otp functions in the adult brain. We have therefore established a C57bl/6 mouse line harboring a “floxed” exon 2 of the *Otp* (*Otp^f/f^*). This mouse line enabled a conditional knockout (cKO) of the Otp gene according to the spatiotemporal expression of the Cre-recombinase employed. Temporal *Otp* deletion was achieved by using Cre fused to the synthetic estrogen receptor ERT2, which requires the introduction of the synthetic ligand tamoxifen to function (20). Spatial expression was restricted to the forebrain using the CamKIIa promoter (21). Crossing the *Otp^f/f^*with *the CaMKIICreERT2* line generated the experimental groups where *Otp^f/f^* mice served as controls, and littermates carrying an additional allele of *CaMKIICreERT2* served as the Otp-cKO mice. At the age of 8 weeks, all mice received tamoxifen by gavage during four consecutive days, which allowed specific induction of Otp ablation in the adult forebrain (Fig. 1A). Depletion of *Otp* mRNA was validated by qPCR of hypothalamic tissue (Fig. 1B). Otp antibody staining, following tamoxifen administration, indicated complete loss of protein translation in the PVN, and the arcuate nucleus of the hypothalamus (Arc) in Otp cKO as compared with control mice (Fig. 1C). These results validate that the tamoxifen-induced deletion of *Otp* exon abolishes expression of Otp transcripts and protein.

**Figure 1.**
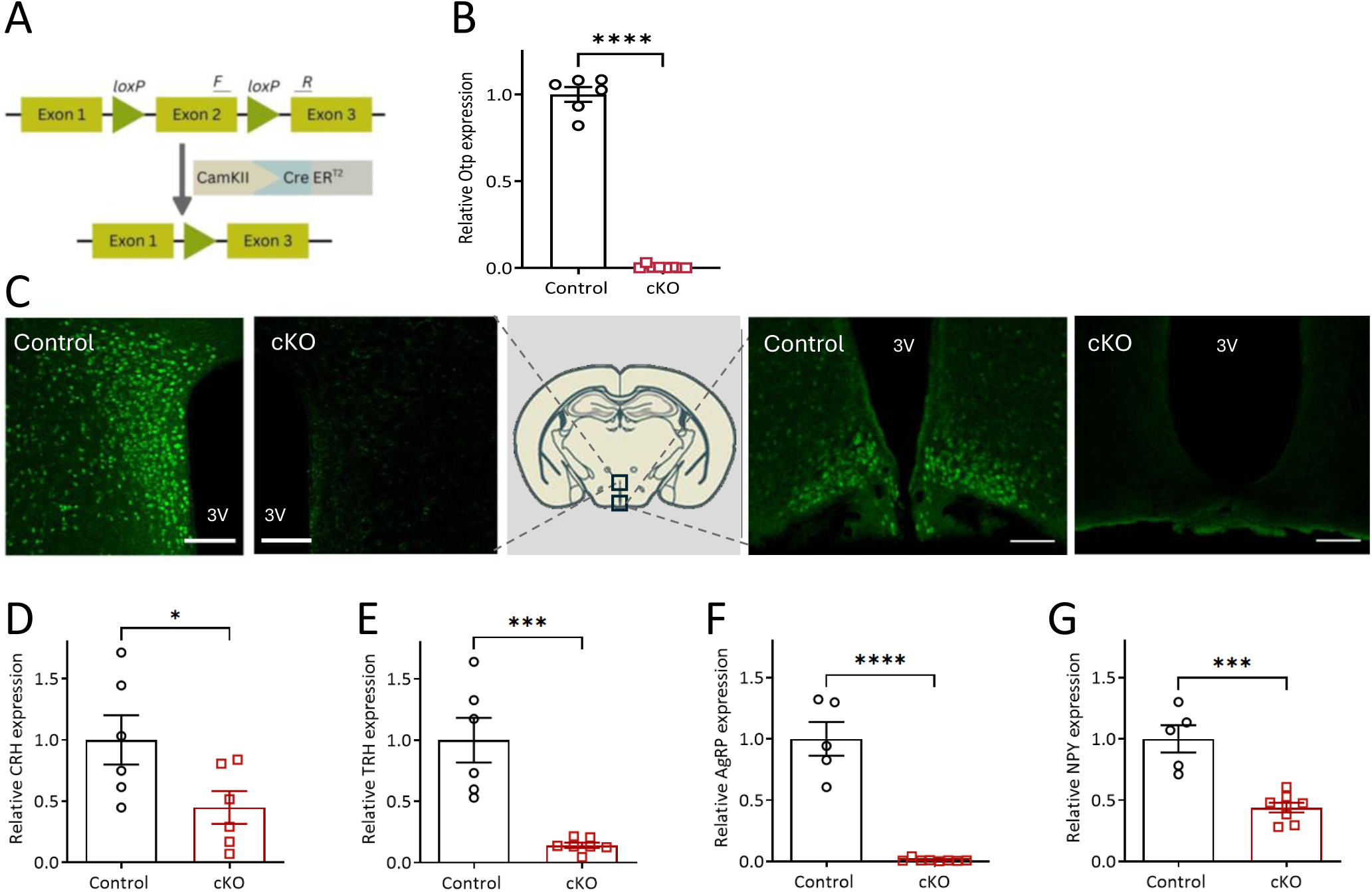
Inducible deletion of Otp exon 2 results in hypothalamic Otp ablation. A. Schematic representation of the cassette used for generating the Otpf/f mice (control) and mice lacking Otp in CamKII-expressing neurons (cKO). B. Relative hypothalamic Otp expression in control and cKO mice (n=6). C. Immunohistochemical staining for Otp using an anti-Otp antibody in the PVN (left) and Arc (right) of control and Otp cKO mice. Scale bar indicates 100μm Relative hypothalamic expression of CRH (D), TRH (E), AgRP (F), and NPY (G) in control and cKO mice (n=6-8). 3V, third ventricle; PVN, Paraventricular nucleus of the hypothalamus; Arc, Arcuate nucleus; AgRP, agouti-related peptide; CRH, corticotropin-releasing hormone; NPY, neuropeptide Y; TRH, thyrotropin-releasing hormone.

### Mice lacking Otp have altered neuropeptide expression and show deficits in hormonal and behavioral adaptation to stress

The *Otp* gene is a downstream target gene of the homeobox transcription factor Dlx1/2 and functions in the proper differentiation of several neurohormone-secreting nuclei and generation of specific neuronal types, among them are the hunger-inducing neurons expressing Aguti-related protein (AgRP) and the SST-expressing neurons (22)(23,24). Given this and the high expression of Otp in the PVN and the Arc, we measured the expression levels of key neuropeptides in these regions in the Otp cKO mice. We found that loss of Otp expression in adulthood led to reduced *corticotropin-releasing hormone* (*CRH*), *TRH*, *AgRP*, and *neuropeptide Y* (NPY) expression (Fig. 1D-G).

During zebrafish development, Otp is required for the differentiation of a variety of neuropeptidergic neurons in the preoptic region, which is the teleost equivalent of the PVN in mice (11,25). We have previously suggested Otp as a component of the homeostatic hypothalamus-pituitary-adrenal (HPA) axis activity, and more specifically, the adaptive response to stress that is associated with its activation (15,16). To test this point, we measured the levels of the stress hormone corticosterone following conditional deletion of *Otp* in the adult forebrain. Basal nocturnal and diurnal levels of plasma corticosterone were not affected by the loss of forebrain Otp (Fig. 2A). Next, stress-induced corticosterone levels were further assessed using acute restraint stress, a paradigm known to induce both psychological and physical stress, leading to a wide range of behavioral and physical alterations, including activation of the hypothalamus to initiate the stress response (26,27). While the Otp cKO mice displayed reduced *CRH* expression (Fig. 1D), they displayed a heightened physiological response to acute stress, as indicated by significantly higher corticosterone levels at 15, 30, and 120 minutes following the restraint initiation (Fig. 2B). Therefore, additional assessments of the behavioral responses to acute stress manipulation were performed. Anxiety-like behaviors utilizing the open-field assay indicated no significant differences between the genotypes under both naïve and stress conditions (Supplementary Fig. 1A,B). Next, assessments of anxiety-like and compulsive-like behaviors were performed using the marble burying test. This test assesses mice’s digging behavior and their tendency to bury unfamiliar, potentially aversive objects in their bedding. Thus, this test is an ethologically relevant assessment of murine stress-coping behaviors in response to potential threats (28). This analysis showed that the Otp cKO mice buried a similar number of marbles under basal conditions (Fig. 2C, left panel), but they displayed an altered sensitivity to stress by burying fewer marbles following acute restraint stress (Fig. 2C, right panel). While the observed stress-induced reduction in marble burying may be interpreted as a reduced anxiety, it corresponds with the Otp cKO mice’s increased corticosterone levels, suggesting a passive stress-coping strategy, akin to not responding in learned-helplessness assays.

**Figure 2.**
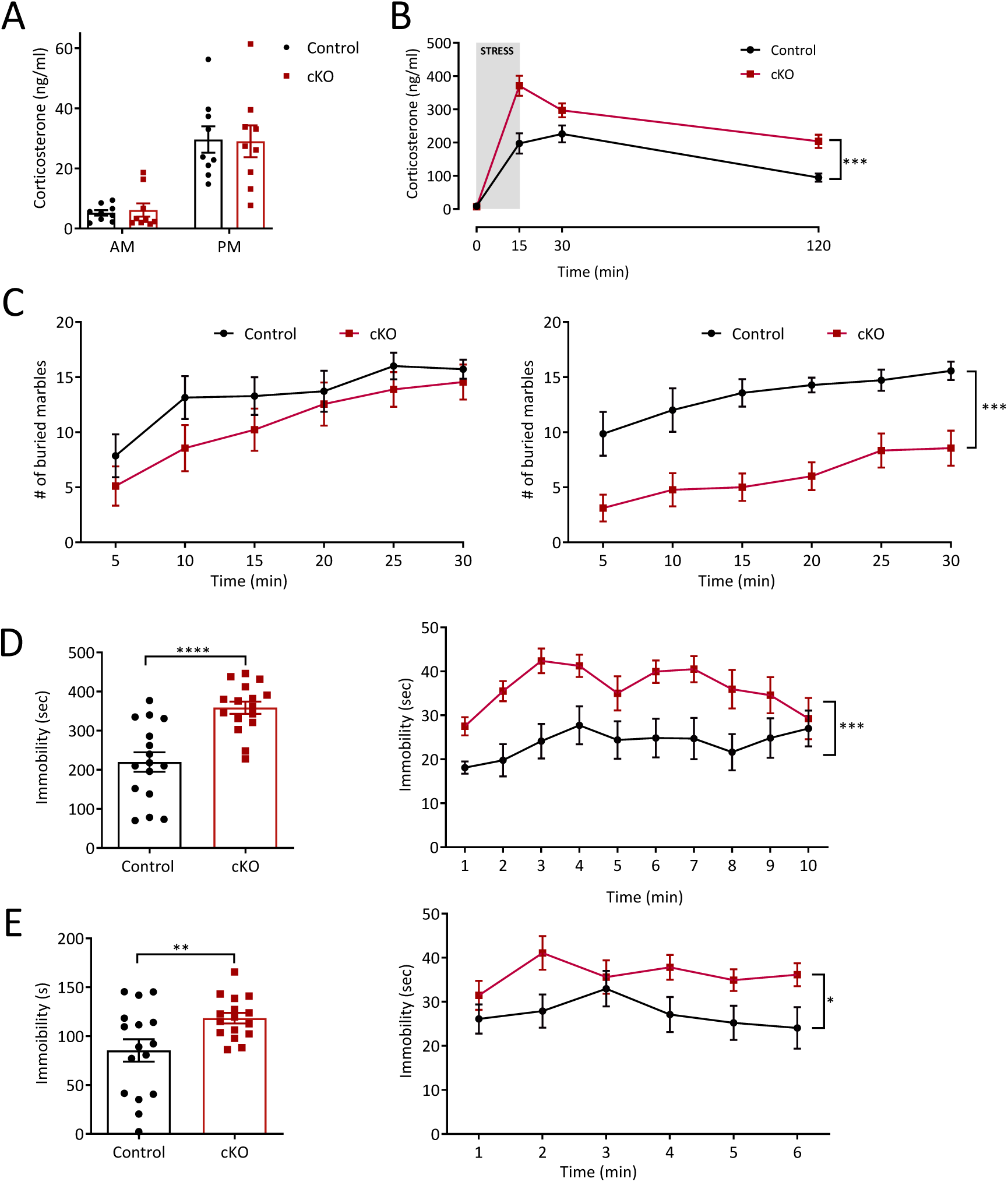
Otp loss in adulthood leads to an increased stress sensitivity. A. Diurnal and nocturnal corticosterone levels (n = 9). B. Restraint stress-induced corticosterone levels (n = 10-11). C. Marble buried under basal state (left) and following 30 minutes of restraint stress (right), shown in 5-minute time bins (n = 7-9). D. Tail suspension total immobility (left) and activity in 1-minute time bins (right, n = 16). E. Forced swim test total immobility as mean (left) and divided into 1-minute time bins (right, n = 16). Data are shown as the mean ± SEM. *p<0.05, **p<0.01, ***p<0.001, ****p<0.0001.

Since increased stress sensitivity and abnormal adaptation to stress are hallmarks of depression and stress-related pathologies, including depression (29), we further assessed passive stress-coping as an indicator of depressive-like behaviors. To this end, we utilized two behavioral assays: the tail suspension test (TST) and the forced-swim test (FST). Both tests are based on measuring the duration of immobility while mice are exposed to an inescapable situation. In mice, immobility in response to uncontrollable stress in these assays is considered a ‘passive’ coping strategy and signifies a behavioral manifestation of despair that can be attenuated by antidepressant medications (30). Compared to control mice, Otp cKO mice displayed increased immobility overall (left panels) and throughout (right panels) the test duration in both TST (Fig. 2D) and FST (Fig. 2E) analyses. To confirm that the immobility phenotype was due to depression-like behavior rather than motor retardation, mice’s motor functions were assessed using the rotarod assay, which assesses balance, coordination, and motion planning (31) as well as using the treadmill exhaustion test. Otp cKO mice did not differ from Control mice in these motor tests, indicating that the loss of Otp in adulthood on motor functions (Supplementary Fig. 1C,D)(31). These results further support a critical role for hypothalamic Otp in the regulation of adaptive coping responses in adulthood.

### Otp loss in adulthood leads to a complex metabolic phenotype

Our current findings demonstrate that Otp loss in adulthood results in reduced expression of neuropeptides that favor energy conservation as AgRP and NPY, while also leading to a reduction in TRH expression, a neuropeptide with an opposing action on energy balance. These key hypothalamic neuropeptides modulate food intake, energy expenditure, and thermoregulation. Moreover, Otp expression in AgRP neurons is induced by fasting and repressed by leptin, a phenotype of orexigenic nature (32). Therefore, we examined neuronal overlap between Otp and metabolic neuropeptides in the adult brain. Using AgRP and SST reporter mice, and immunostaining for Otp and ACTH, we visualized and quantified the expression and co-localization of Otp protein in these major neuroendocrine neurons of the Arc nucleus. We found that Otp is expressed in most of the orexigenic AgRP neurons (91% ± 7%, n=3, Fig. 3A,D), and in a small subset of the adult anorexigenic POMC neurons (identified by staining for ACTH (6.5% ± 3%, n=3, Fig. 3B,D). In addition, Otp is expressed in about half of the SST-expressing neurons (52% ± 14%, n=2, Fig. 3C,D), suggesting a role for Otp in growth hormone regulation. The expression of Otp in the adult Arc neurons further supports its involvement in energy expenditure and adaptive thermogenesis in adult mice. To examine Otp’s physiological effect, we inserted a temperature transponder (IPTT-300) and measured the subcutaneous temperature of the mice. The initial body temperature was similar among all mice, regardless of the presence of the ERT2-cre transgene (Fig. 4A). Two weeks after the cre induction by Tamoxifen administration (shaded grey bars) and onward, the body temperature of mice lacking Otp was significantly reduced, and it remained lower thereafter; indicated by a significant (p=0.012) genotype x time interaction. This was accompanied by reduced blood T4 levels (Fig. 4B) and elevated blood cholesterol levels (Fig. 4C), all of which are hallmarks of an altered hypothalamic-pituitary thyroid (HPT) axis. Metabolic alteration was also observed in fasting glucose levels, which were significantly higher for Otp cKO mice (Fig. 4D inset). However, the mice’s ability to handle a glucose load was intact (Fig. 4D). In addition, the body weight of Otp cKO mice was comparable to that of control mice (Fig. 4E). However, body composition measurement revealed a higher percentage of fat and a lower percentage of lean mass in the cKO mice, consistent with altered thyroid hormone function (Fig. 4F). Indirect calorimetry showed no differences in heat production or food intake (Fig. 4G,H). This result was somewhat unexpected, given the altered body composition and the observed tendency of the cKO mice to be less active (Fig. 4I, p=0.061). Nevertheless, it aligns with the results of a recently described mouse model with reduced PVN Otp expression (18). Water intake was significantly higher in Otp cKO mice (Fig. 4J), probably due to altered AVP expression, which is accompanied by significantly lower urine osmolality (Fig. 4K, significant, p=0.032, interaction: genotype x treatment). Despite the lower basal temperature, Otp cKO mice defended their body temperature comparably to controls when exposed to a cold environment, maintaining the basal difference (Supplementary Fig. 2A) and exhibiting a similar induction of uncoupling protein 1 (UCP1) expression (Supplementary Fig. 2B).

**Figure 3.**
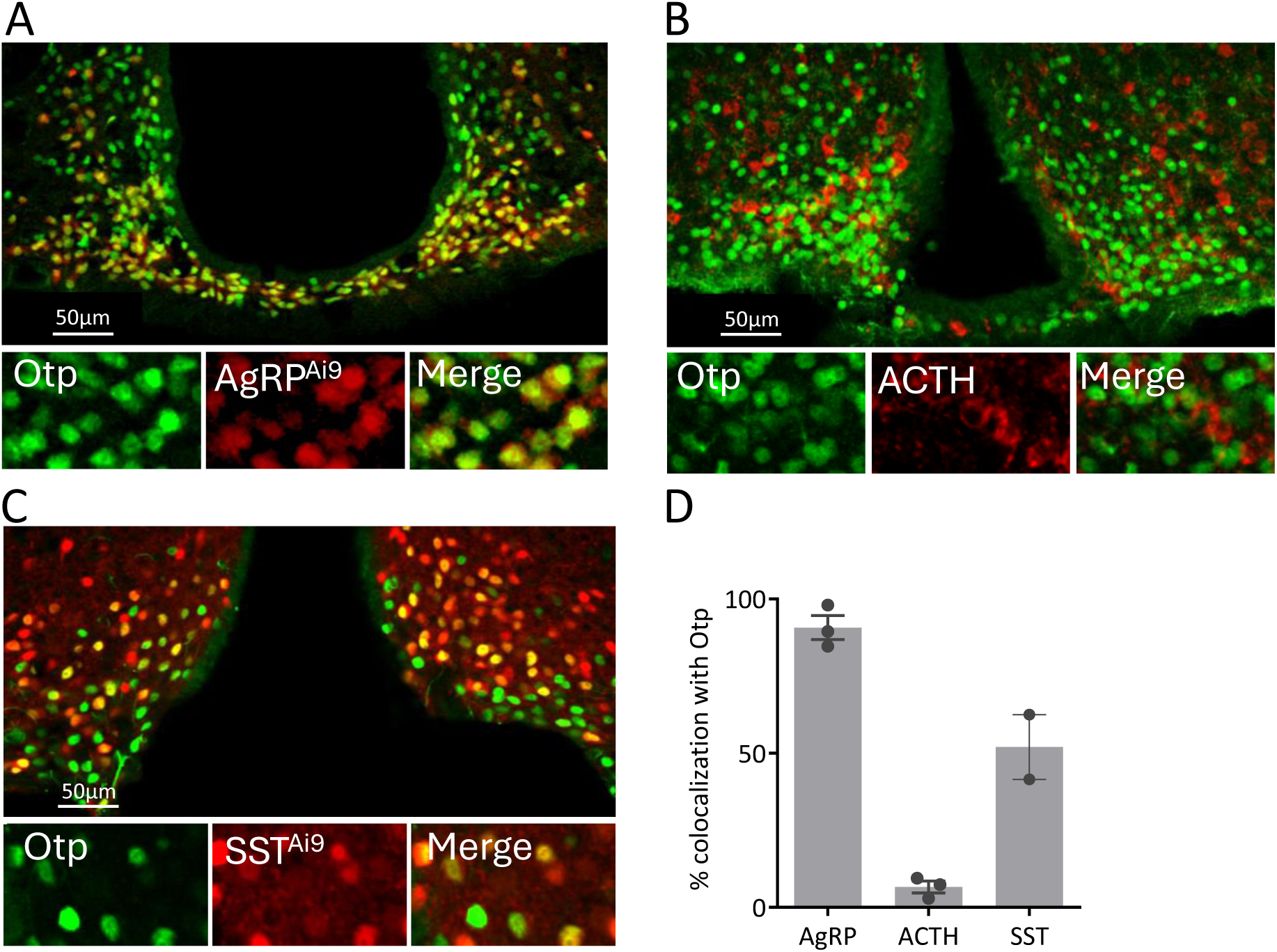
Otp is expressed in several neural populations in the arcuate nucleus. A. Immunodetection of Otp in the Arc of AgRP reporter mouse. B. Immunodetection of Otp and POMC precursor, ACTH, in the Arc of colchicine-injected mice. C. Immunodetection of Otp in the Arc of SST reporter mouse. D. Percentage of Otp-positive subpopulation out of total AgRP, ACTH, or SST expressing neurons (n=2-3).

**Figure 4.**
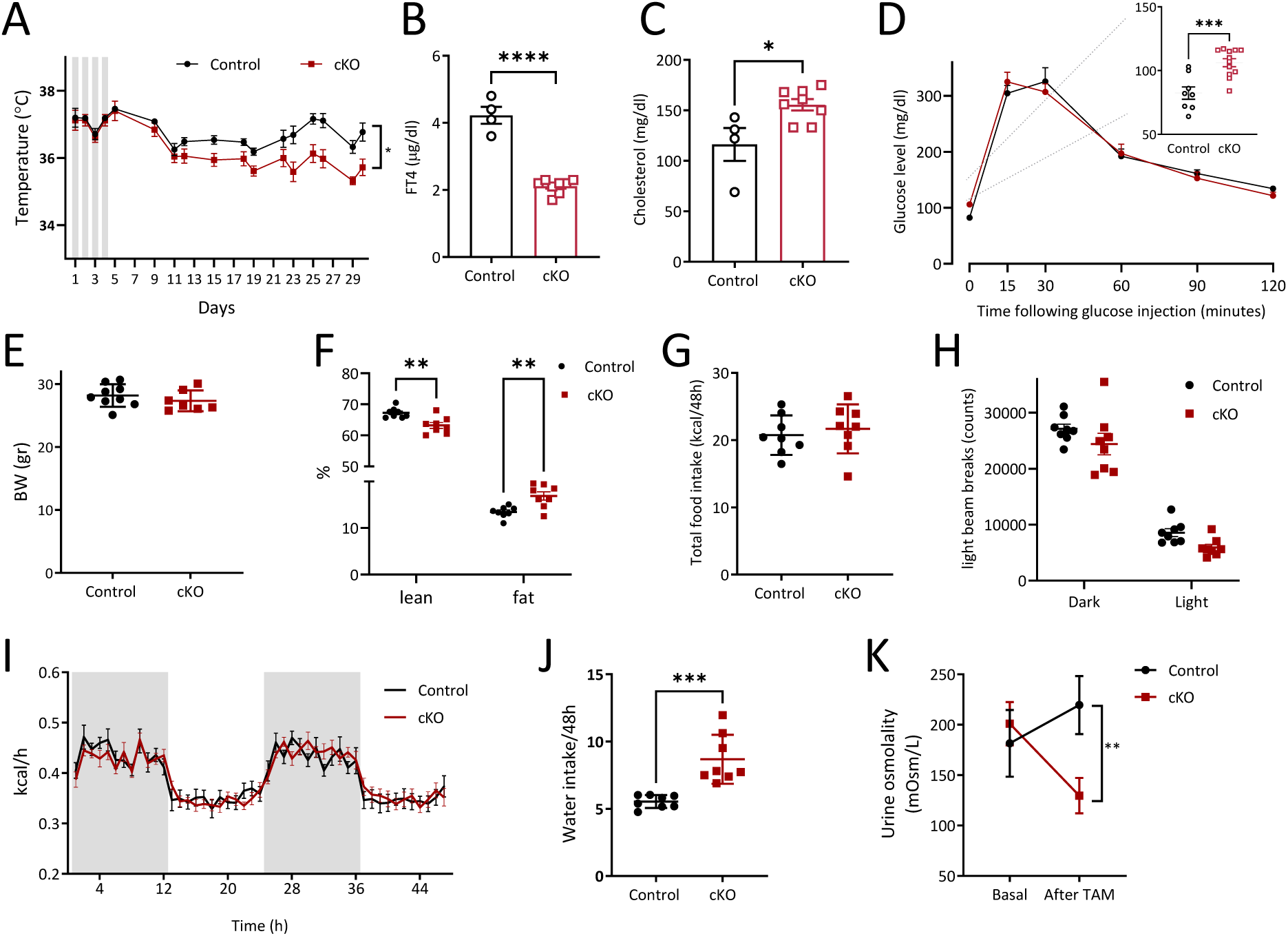
Otp loss in adulthood leads to an altered HPT axis and to a complex metabolic phenotype. A. Subcutaneous temperature of the mice measured from the first day of tamoxifen administration (depicted by gray bars, n = 8,9). B, C. Blood free-T4 and Cholesterol levels (n = 4-8). D. Glucose tolerance test and fasting glucose in inset (n=8,12). E. Body weight and F. body composition upon entering the metabolic cages (n=8). G. Food intake during 48h. H. Heat production kinetics. I. Averaged locomotion during each light phase. J. Water intake during 48h. K. Urine osmolality, before and three weeks after tamoxifen administration (n=6,8). Data are shown as the mean ± SEM. *p<0.05, **p<0.01, ***p<0.001, ****p<0.0001.

Overall, these data demonstrate that in adulthood, Otp is essential for maintaining thyroid hormone functions and their associated metabolic roles; therefore, its absence results in a complex metabolic phenotype.

### Otp loss in adulthood leads to reduced responsiveness to ghrelin, which is not due to the death of AgRP neurons

Otp is required for the generating of AgRP neurons and their continuous differentiation after the fate-cell determination (22). The dramatic reduction in AgRP expression, observed following deletion of Otp in adulthood, could be due to various reasons and may result from Otp’s effects on cell viability or the peptide mRNA levels. Given the dramatic orexigenic effect of ghrelin signaling in AgRP neurons (33), we evaluated the feeding response to ghrelin administration to Otp cKO mice. We intraperitoneally injected the mice with ghrelin during the light phase, when feeding is minimal, and measured their food intake 30 and 60 minutes after the injection. As compared with control conditions (saline injection), food intake in response to ghrelin injection was significantly lower in Otp cKO mice compared to control mice; this difference was observed already by 30 minutes following the injection and the difference between the genotypes was substantially increased at 60 minutes post injection (Fig. 5A, significant, p=0.002, 3-way interaction: genotype x time x treatment). We further examined the medio-basal hypothalamic expression of genes enriched in AgRP neurons (34), including the ghrelin receptor (*growth hormone secretagogue receptor (GHSR)), Serpina3n, activin A receptor type 1C (Acvr1c),* and *5-hydroxytryptamine receptor 1B (5-HT1B)*. We did not find a significant change in the expression of these genes (Fig. 5B), suggesting that AgRP neurons remain viable following Otp deletion. To further investigate whether Otp deletion induces cell death of part of the AgRP neurons, we evaluated the presence of apoptotic markers. To this end, mice were sacrificed one week following the first tamoxifen administration, and brain sections were stained for Galectin-3, Glial Fibrillary Acidic Protein (GFAP), and Cleaved Caspase 3. A complete depletion of Otp protein in cKO mice was reached one week following tamoxifen administration. However, no difference in the staining pattern was observed for both Galectin-3 and GFAP (Fig. 5C), and no staining of Cleaved Caspase 3 was detected (not shown).

**Figure 5.**
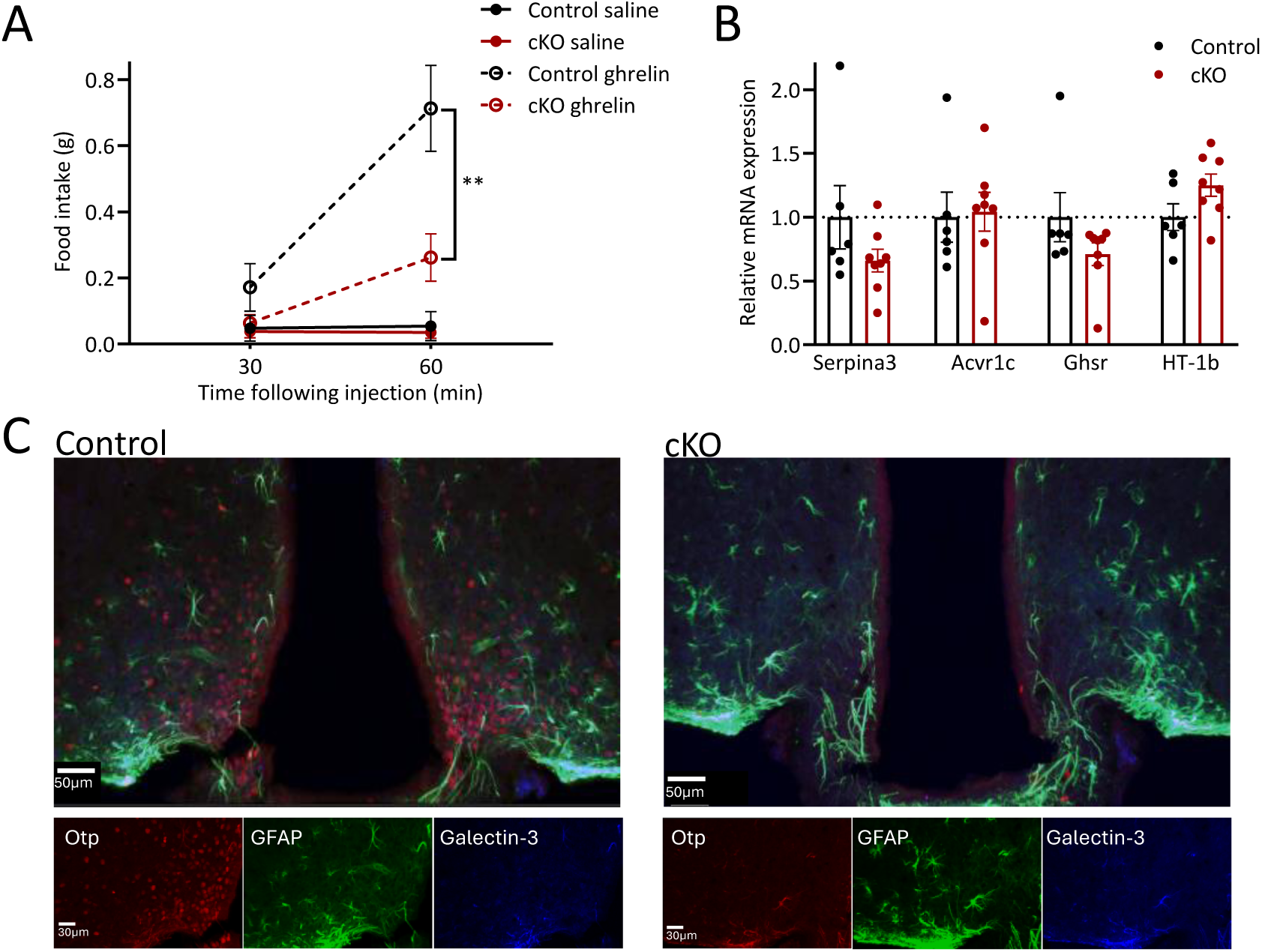
Otp loss in adulthood leads to reduced responsiveness to ghrelin, which is not due to loss of AgRP neurons. A. Food intake following saline or ghrelin injection (n=5) B. MBH expressions of Serpin3a, Acrvr1c, GHSR and HT1B (n = 6-8) C. Immunostaining for Otp, GFAP, and Galectin-3 one week after Tamoxifen administration to control or cKO mice. Data are shown as the mean ± SEM. **p<0.01.

To summarize, these results indicate that conditional depletion of Otp protein in the adult Arc nucleus leads to loss of *AgRP* expression, which affects the activity of the neurons that previously expressed AgRP without affecting their viability.

## Discussion

Otp is an evolutionary conserved transcription factor that has a critical role in hypothalamic neuroendocrine development. The expression of Otp in specific neurons of the adult hypothalamus of fish and mammals indicates its possible involvement in regulating physiological stress response and energy balance. To examine this conjecture, it was necessary to uncouple the neurodevelopmental effects of Otp from its presumed role in adult physiological function. Using a conditional forebrain-specific mouse model, we now demonstrate that in addition to its established role in hypothalamic neuronal differentiation, Otp serves as a master regulator of neuroendocrine functions in adults.

In our Otp cKO model, Otp is deleted during adulthood in several forebrain regions, including the PVN and the Arc. These regions control various homeostatic functions, and therefore, the described multifaceted phenotype may be attributed to the deletion of Otp in any of these areas, or to their interactive combination. Indeed, our findings highlight the pleiotropic roles of Otp in the adult brain.

The HPA axis is a prominent arm of the stress response, and together with the HPT axis, CRH and TRH modulate sympathetic outflow and affect energy balance (35). Here, we show that the expression of both upstream players, CRH and TRH, is reduced in Otp cKO mice. This led to an increase in stress reactivity, despite the lower CRH expression, as well as to central hypothyroidism with its related behavioral and metabolic side effects. Loss of Otp in adulthood led to a heightened corticosterone response to stress, coupled with increased depressive-like behaviors. HPA axis overactivation is a prominent and well-documented neuroendocrine abnormality in depression (36). The expression of CRH, as well as alteration in cortisol secretion, is commonly found to be dysregulated in various human neuropsychiatric disorders, such as depression, anxiety disorders, and post-traumatic stress disorder (37). Dysfunction of the HPT axis is another known neuroendocrine abnormality in depression, which is altered in a substantial proportion of depressed patients (38,39). Like the HPA axis, dysregulation of the HPT axis correlates with depression; specifically, thyroid hypofunction was shown to lead to long-term sub-clinical levels of depression, previously termed dysthymia, and to decreased levels of psychomotor activity (40). Moreover, the combination of HPA and HPT axes perturbations has a synergic contribution to the severity of neuropsychiatric disorders (41). The dysregulated HPA and HPT axes may, at least in part, underlie the shift towards passive coping behaviors observed following the loss of Otp in adulthood in the FST and TST assays. The decrease in NPY expression observed in our mouse model may also contribute to the behavioral phenotype of Otp cKO mice. NPY plays a role as a resilience-promoting factor, and its reduced function is linked with exaggerated stress and anxiety responses (42,43). Accordingly, chronic stress was shown to reduce AgRP expression and to alter firing properties of AgRP neurons, while chronic inhibition of AgRP neurons increases anxiety-like and depressive-like behaviors (44,45).

The Ghrelin hypo-responsiveness is also in line with the increased anxiety-like and depressive-like behaviors. Ghrelin was implicated in regulating stress responses in laboratory rodents and was suggested as a potential therapeutic target in humans who suffer from stress-related psychopathologies, including depression (46,47).

Beyond the behavioral outcomes, dysregulated HPA and HPT axes can alter energy balance by affecting food intake, energy expenditure, and autonomic outflow. This can be mediated both via central mechanisms and by the circulating effectors-corticosterone and T3/T4 (48).

Developmental mouse models with alterations in Otp are obese and hyperphagic, a phenotype that may be, at least in part, attributed to hypothyroidism (17). In the present study, we studied the postnatal metabolic function of Otp. This can be compared to the metabolic phenotype of a recent study describing mice in which Otp expression in the PVN was reduced in adulthood by employing a Cre-expressing viral vector (18). In both models, Otp expression was manipulated in adulthood, allowing proper brain development and neural wiring. In both models, reduced Otp expression in adulthood led to a higher fat percentage, with intact heat production and locomotor activity. However, as the developmental Otp KO models, the PVN-specific Otp depletion had positive energy balance resulting in hyperphagia and obesity that were not present in the Otp cKO we report herein. The main difference between the two is that in our model, Otp expression was reduced in both PVN and the arcuate nucleus, and this resulted in a dramatic reduction in AgRP and NPY expression, leading to reduced responsiveness to hunger signals and thus halting body weight gain. Increased water intake in our Otp cKO mice may also contribute to the satiation perception. Others have reported that AgRP neuron ablation in adulthood has not altered body weight and food intake (49). However, in the present study, neurons were not ablated, but the expression of their signature neuropeptides was reduced. Reduction in AgRP expression was reported to increase heat production without affecting food intake, leading to a reduction in body weight (50). We propose that in our current Otp cKO mice, the effects of Otp loss in both the arcuate nucleus and the PVN may counterbalance each other. This may occur since Otp loss influences the expression of both orexigenic and anorexigenic neuropeptides, which mask obesity and hyperphagia phenotypes, which are observed in the PVN-specific mouse model. This phenotype positions Otp in the hypothalamic as a potential candidate in combating pathological eating disorders and central hypothyroidism, such as seen, for example, in Prader-Willi syndrome (51,52).

We previously showed that Otp regulates the balance of neuropeptide production in zebrafish (2,16). We now suggest that Otp has an opposing role in neighboring hypothalamic nuclei. Thus, Its action cannot be classified as having simple anxiogenic or anxiolytic-like downstream effects and cannot be classified as having a role only in energy conservation or energy dissipation.

Collectively, the data presented herein indicate that, in adulthood, Otp is a key regulator of several neuroendocrine axes in the complex and interrelated neuronal networks that underlie homeostatic stress response and energy balance. Malfunctions of these systems may contribute to obesity, depression, and various other stress-related neuropsychiatric disorders and therefore warrant additional research.

## Methods

### Animals

All animals used for the experiment were naïve adult males, housed and handled in a specific pathogen-free temperature-controlled (22°C ±1) mouse facility on a 12/12h light/dark cycle (lights on from 6:00–18:00 h, or 22:00-10:00 in a reverse cycle), with food and water given ad libitum according to institutional guidelines. The Institutional Animal Care and Use Committee of the Weizmann Institute of Science approved all experimental protocols. The following mouse lines were obtained from Jackson Laboratory, Bar Harbor, ME, USA, and used in this study: AgRP-Cre, stock number 012899; CamKII Cre-ERT2, Stock number 012362; Floxed Ai9, (Rosa26-CAG-lsl-tdtomato) stock number 007909. Somatostatin-cre mouse line was kindly provided by the Lample lab, Weizmann Institute of Science.

### Generation of Otp-cKO mice

Mutant ES cells containing floxed-Otp allele Otptm1a^(EUCOMM)Wtsi^ (orthopedia homeobox; targeted mutation 1a, Wellcome Trust Sanger Institute) were obtained from EUCOMM (The European Conditional Mouse Mutagenesis Program, by Wellcome Trust Sanger Institute, Cambridgeshire, UK).

For the floxed-Otp allele, the L1L2-Bact-P cassette was inserted at position 94876664 of Chromosome 13 upstream of the critical exon, which contains the homeobox domain (Build GRCm38). This cassette includes an FRT site followed by a lacZ sequence, a loxP site followed by neomycin under the control of the human beta-actin promoter, SV40 polyA, a second FRT site, and a second loxP site. A third loxP site is inserted downstream of the targeted exon at position 94877768. The Homeobox-containing exon is thus flanked by two loxP sites. By introducing flp recombinase in mice carrying this allele, a “conditional ready” (floxed) allele was created, while subsequent cre expression results in a knockout mouse. The mutant ES cells were used to generate chimeric mice by blastocyst injection. Germ-line transmission of the modified *Otp* allele (*Otp^loxP^*) was confirmed in offspring from male chimeras bred to wild-type C57BL6 mice. Finally, the *frt* flanked selection cassette was removed by breeding with *Deleter-Flp* mice. To achieve spatiotemporally controlled somatic mutagenesis, Cre-recombinase was introduced by crossing the Otp-floxed mice with CaMKCreERT2 transgenic mice that express Cre recombinase under the CamKIIa promoter, known to be widely active in forebrain neurons. In this construct, a mutated form of estrogen ligand-binding domain (ERT2) that binds to synthetic antagonists such as tamoxifen but not to circulating estrogens, was fused to the Cre to create the CreERT2, which requires the presence of tamoxifen for Cre recombinase activity (Fig. 2A). PCR genotyping was performed on genomic DNA extracted from tail biopsies. Otp-loxP forward 5’-atggatgtgactccgtttccc-3’ and reverse 5’-gtttgagtactccaccccgc-3’ primers yield a 1476-bp product from the WT gene and a 1613-bp product from the floxed Otp allele. Cre forward 5’-ggttctccgtttgcactcagga −3’ and reverse 5’-gcttgcaggtacaggaggtag −3’ primers yield a 375-bp product that is specific to the transgene allele.

For genomic validation of Otps’ deletion, PCR was performed using primers flanking the loxP sites, which flank Exon 2. A processed product of 547b was observed in Otp^f/f^; CaMKIICreERT2^-/+^, but not in Otp^f/f^; CaMKIICreERT2^-/-^ mice. This band was extracted from the gel and sent for sequencing, which revealed a genomic deletion of exon 2 in the Otp gene and the remains of one Frt and one loxP site, as expected.

### Tamoxifen administration

Tamoxifen (Sigma) was dissolved in ethanol at 37.5 mg/ml and kept at −20°C. Before usage, corn oil (Sigma) was added at the same volume as ethanol, and ethanol was evaporated using Concentrator 5301 (Eppendorf). Tamoxifen was administered to the mice via gavage at a 150 mg/kg dose for four consecutive days. To induce Otp deletion, and to avoid bias due to tamoxifen-related effects (53), all mice received tamoxifen. At the age of 8-10 weeks, mice of both Otp^f/f^; CaMKIICreERT2**^-/+^** (cKO) and Otp^f/f^; CaMKIICreERT2**^-/-^** (control) genotypes were administered with tamoxifen. Unless specified otherwise, behavioral or molecular assessment occurred at least two weeks following tamoxifen administration.

### Intraventricular injection of colchicine

For immunostaining secreted neuropeptides, mice were injected with colchicine (2μg) into the lateral ventricles (AP −0.22 mm; ML +0.95 mm; DV −2.2 mm). Mice were sacrificed when locomotor symptoms were observed.

### Restraint stress

Restraint was applied by placing mice in ventilated 50-ml polypropylene Falcon tubes for 30 min, followed by a 15-minute recovery in a new cage.

### Forced-swim test

Mice were forced to swim in a bucket (20 cm Ø) filled with water to a depth of 30 cm at 25± 2°C, while a camera recorded the session. The test consisted of two sessions on two consecutive days: 15 min on day 1 as ‘induction’, followed by a 5 min ‘trial’ performed 24 hours later. Immobility was used as an index of learned helplessness (behavioral despair), indicative of a depressive-like state. Immobility was automatically scored using EthoVision XT (Noldus) software, measuring the percent of change in pixels; immobility was defined as <1% change for at least 1 second.

### Tail suspension test

Mice’s behavioral despair was assessed using the tail suspension system (TSE-systems; Berlin, Germany). Mice were suspended upside down for 6 minutes by tapping the tip of their tail to a metal chain connected to a sensitive motion sensor. The resulting escape-oriented behaviors were quantified by measuring the duration of each mouse’s struggle and the magnitude of the swings generated by this movement.

### Marble-burying assay

The mice’s responses to novel/unfamiliar objects (marbles) were assessed among naïve (unstressed) mice and mice exposed to an acute restraint stress challenge of 30 min, followed by a 15-minute recovery. Mice were placed for 30 min in an arena (25 cm × 25 cm × 30 cm) filled with 5 cm of wood chip bedding and 20 evenly spaced marbles. The assay was video recorded, and the number of marbles buried was counted every 5 minutes.

### Open field test

The open field apparatus (TSE-systems, Berlin, Germany) consists of a white plexiglass box (50 cm × 50 cm × height 40 cm), illuminated with 120 lux. Each mouse was placed in the corner of the apparatus to initiate a 10-minute test session. A camera (Eneo, Rödermark, Germany; Model: VK1316S) mounted above the apparatus transmitted images of the mice, which were analyzed by TSE-systems’ VideoMot2 software. The following indices were recorded and subsequently analyzed: time spent in the center, number of visits to the center, and total distance traveled. More frequent or longer times spent exploring the arena or in the center of the arena are indicative of anxiety.

### Treadmill exhaustion assay

Mice were acclimated to treadmill running (Panlab Harvard Apparatus, Barcelona, Spain) for 2 days before the endurance exhaustion test. The pre-tests included 10 min at 10 cm/s, while the test was composed of: phase I – 10 min at 10 cm/s, phase II – increases by 0.02 cm/s every min for 10 min; Phase III - 10 min of maximal speed of 0.35 cm/s. Mice ran until exhaustion, defined as 20 cumulative seconds of no running. Time to exhaustion was recorded and compared.

### Rotarod performance test

Mice’s motor coordination and balance were tested using the RotaRod system (San Diego Inst.). Mice were placed on a horizontally oriented, rotating cylinder (rod) suspended above a cage floor with the speed increasing linearly from 0 to 40 rpm over a 4 min. The protocol included a habituation day, composed of 9 trials in which the mice learned to balance and coordinate their movement on the rotating rod, while mice falling from the rod were immediately placed back on it. Day 2: test session composed of 5 trials with a 2-minute break between trials. Mice’s latency to drop off rotating rods was recorded and served as an index of performance.

### Immunohistological analysis

Animals were anesthetized and transcardially perfused with 4% paraformaldehyde (PFA). Brains were post-fixated with 30% sucrose solution in 4% PFA. Fixed brains were serially sectioned into three sets of 40μm slices. Slices were then incubated in a blocking solution (20% horse serum, 0.05% Triton in phosphate-buffered saline (PBS) for 2 hours to prevent nonspecific binding of the antibody. Next, sections were incubated for 24 hours at room temperature and 24 hours at 4°C with the primary antibody in PBS containing 2% horse serum and 0.2% Triton. Following PBS washes, sections were incubated at room temperature for 1-2 hours with the appropriate secondary antibodies (Jackson Immunoresearch Laboratories Inc., West Grove, PA, USA). Sections were washed with PBS and mounted on slides. Images were captured using a confocal microscope (Zeiss LSM880 and LSM700). For quantification of Otp colocalization, a complete set of Arc slices was images at 2μm intervals. Otp antibody (working concentration 5 ng/ul) was raised against a C-terminal Otp peptide and purified by affinity chromatography as previously described (54). The following antibodies were also used: ACTH (1:1000, DAKO), Caspase-3 (1:50), and GFAP (1:100).

### Brain regions microdissection

For region-specific analysis, brains were microdissected using the Palkovits technique for punch sampling as described (55). Immediately after the decapitation, the brain was removed and placed into a 1mm acryl matrix (Stoelting Co., 51386). Brains were sliced using standard razor blades (GEM, 62-0165) into 2 mm slices. Punches were taken using an 18-G blunted needle and were quickly frozen for further analysis.

### RNA Extraction and Real-Time PCR

mRNA from dissected tissue was extracted using the RNeasy mini or micro kit (QIAGEN). cDNA was synthesized from 0.5–1 mg of total RNA using an oligo(dT)15 primer and SuperScript II reverse transcriptase (RT) (Invitrogen). Samples were then analyzed using the miScript SYBR Green PCR kit (QIAGEN), according to the manufacturer’s guidelines, in the AB 7500 thermocycler (Applied Biosystems). cDNA products were used as templates for real-time PCR analysis. Quantitative (q) PCR analysis was done using the StepOnePlus system using SYBR Green (Applied Biosystems, Life Technologies Corp., CA, USA). Primer sequences will be provided upon request.

### Plasma glucocorticoid measurement

Tail blood was collected under basal “naïve” conditions, at either the active phase or non-active phase (9 am and 9 pm), and at several time points (15, 30, and 120 minutes) after the initiation of a 15-minute restraint stress. Bleeding was induced by cutting the tip of the tail, and blood (20 μl) was collected by a designated pipette into an Eppendorf tube containing 5μl of EDTA 0.5M, pH 8.0, and kept on ice. Plasma was separated by centrifugation and kept at −20°C until use. Corticosterone levels were determined by radioimmunoassay (RIA) using a corticosterone double-antibody ^125^I RIA kit (MP Biomedicals Inc.) according to the manufacturers’ instructions.

### Serum total T4 and cholesterol measurements

100μl of blood was drawn from the mice’s cheek and analyzed using Vetscan VS2 Chemistry Analyzer (Zoetis, NJ, USA) and the T4/Cholesterol Profile.

### Metabolic Cages Analysis and Body Composition

Indirect calorimetry, food, water intake, and locomotor activity were measured using the PhenoMaster system (TSE-Systems, Berlin, Germany). Data was collected after 48 hours of adaptation in acclimated singly housed mice. Mice’s body composition was assessed using Bruker minispec mq7.5 Live Mice analyzer (Bruker BioSpin, Billerica, MA, USA).

### Temperature measurement and cold challenge test

Mice were implanted with an Implantable Programmable Temperature and Identification Transponder (IPTT-300; Bio Medic Data Systems, Avidity Science, USA) under isoflurane anesthesia. Body temperature data were obtained using a SP-6005 reader. For the cold challenge, mice were singly housed and, two hours afterward, were placed at 4°C. Temperature was measured in 30-minute intervals.

### Urine osmolality

Urine was diluted 1:10 with ddw, and osmolality was measured using a Fiske 210 Micro-Sample Osmometer (FISKE® ASSOCIATES, MA, USA).

### Ghrelin sensitivity test

Ghrelin (3 µg/g) was injected i.p. into fed mice during the light phase, and food intake was measured after 30 and 60 minutes. A similar volume of saline was injected on the previous day as a control.

### Glucose tolerance test

For the glucose tolerance test (GTT), glucose (2g/kg body weight) was injected i.p. following five hours of fasting. Whole venous blood glucose obtained from the tail vein at 0, 15, 30, 60, 90, and 120 min after injection was measured using an automatic glucometer (Abbott).

### Data analysis and statistics

Indices are expressed as means +/- SEM. Statistical analyses were performed using PRISM software (GraphPad Software, Inc., La Jolla, CA, USA). Student’s t-tests and two-way ANOVA (with in-subjects repeated measures) followed by post-hoc Bonferroni corrected t-tests were used as appropriate. * Indicates p<0.05, ** p<0.01, ***p<0.001, ***p<0.0001.

## Author contributions

Conceptualization, M.Ta., Y.K., A.C., and G.L.; Methodology, M.Ta. T.N., M.Ts., B.B., J.Bi., E.R., and Y.K.; Data collection/ Investigation, M.Ta., T.N., B.B., E.R., and Y.K; Data curation and analysis, M.Ta., M.Ts., and Y.K.; Writing-original draft, M.Ta., Y.K., and G.L.; Writing-review and editing, M.Ts, J.Bi., A.C., M.Ta., Y.K., and G.L.; Supervision, Y.K., A.C., and G.L.; Funding acquisition, G.L., and A.C.; Project administration, A.C., G.L., and Y.K.

## Acknowledgement

G.L. is supported by the Israel Science Foundation (#349/21); the Hedda, Alberto, and David Milman Baron Center for Research on the Development of Neural Networks; the Minerva Foundation with funding from the Federal German Ministry for Education and Research, the Wolfe Family Center for Research on Neuroimmunology and Neuromodulation; the Sagol Center for Research on the Aging Brain; the Irene and Jared M. Drescher Center for Research on Mental and Emotional Health; the Azrieli Institute Center for Research on the Development of Innovative Neuro-technologies; and the Maurice and Vivienne Wohl Biology Endowment and Foundation for Higher Education and Culture. G.L. is an incumbent of the Elias Sourasky Professorial Chair. This work was conducted at the Ruhman Family Laboratory for Research on the Neurobiology of Stress and was supported by research grants (to A.C.) from Bruno and Simone Licht; the Leff Family; the Irving B. Harris Fund for New Directions in Brain Research; the Joseph D. Shane Fund for Neurosciences; the Estate of Ethel Lena Levy; the Benoziyo Endowment Fund for the Advancement of Science; the Estate of Hermine Miller; the Estate of Gertrude Buchler; the Estate of Marjorie Plesset; the Estate of Zvia Zeroni; the Estate of Olga Klein Astrachan; the Estate of Gerald Alexander; and the Anita James Rosen Foundation. A.C. is the incumbent of the Vera and John Schwartz Family Professorial Chair in Neurobiology at the Weizmann Institute of Science. M.Ts. is the incumbent of the Carolito Stiftung Research Fellow Chair in Neurodegenerative Diseases. Y.K. is the incumbent of the Sarah and Rolando Uziel Research Associate Chair.

---

**Supplementary Figure 1.**
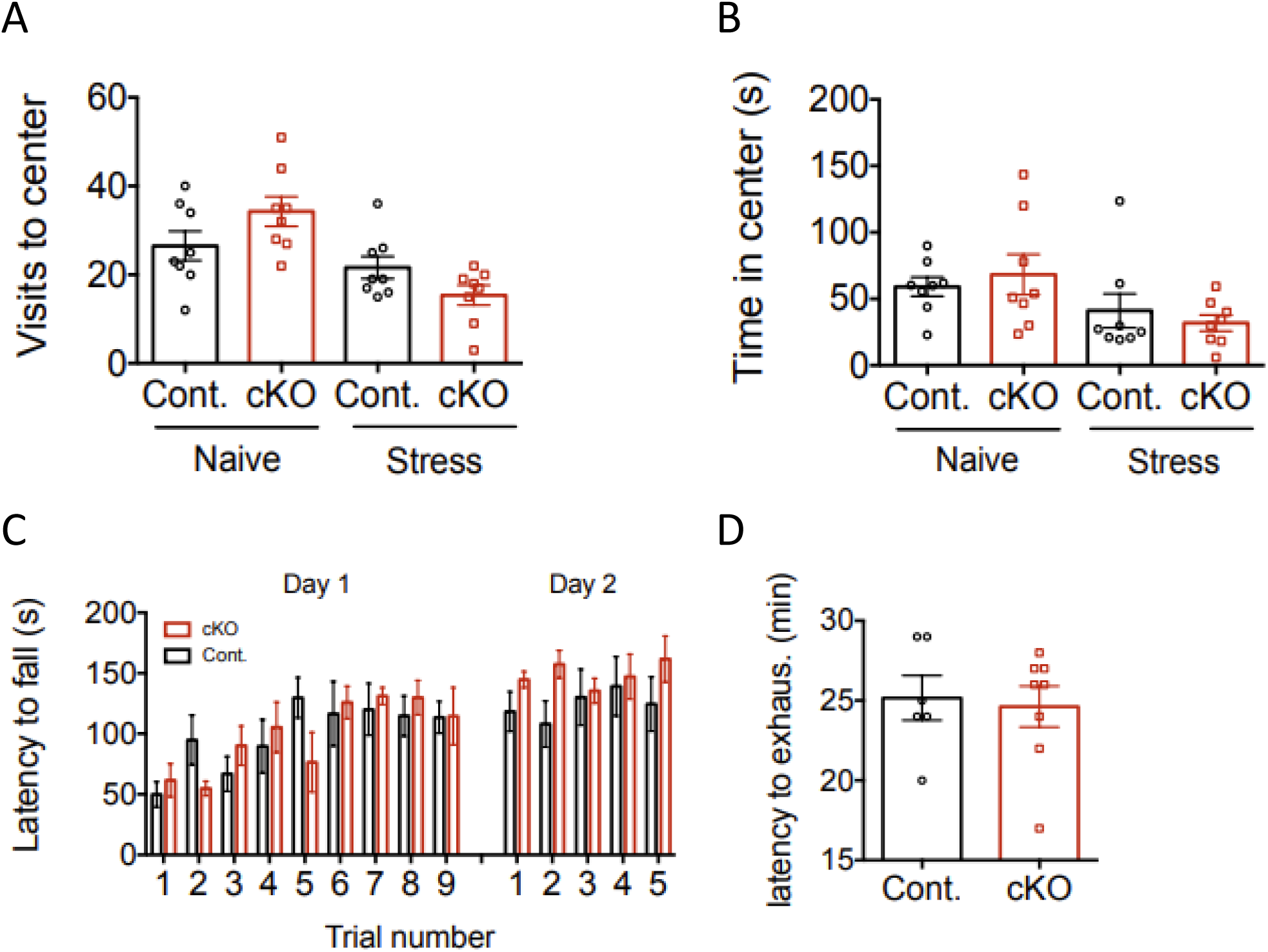
Otp loss in adulthood does not alter anxiety-like behavior or locomotor function. A. Visits to the center of the open-field arena in mice tested under basal conditions or 15 minutes following a 30-minute period of restraint stress (n=8). B. Time spent in the center of the open-field arena in mice tested under basal conditions or 15 minutes following a 30-minute period of restraint stress (n=8). C. Motor coordination and balance assessed by the latency to fall of a rotating rod during 9 training trials on the first day and 5 performance trials on the second day (n=8). D. Endurance assessed by the latency to exhaustion while running on the treadmill (n=6,8).

**Supplementary Figure 2.**
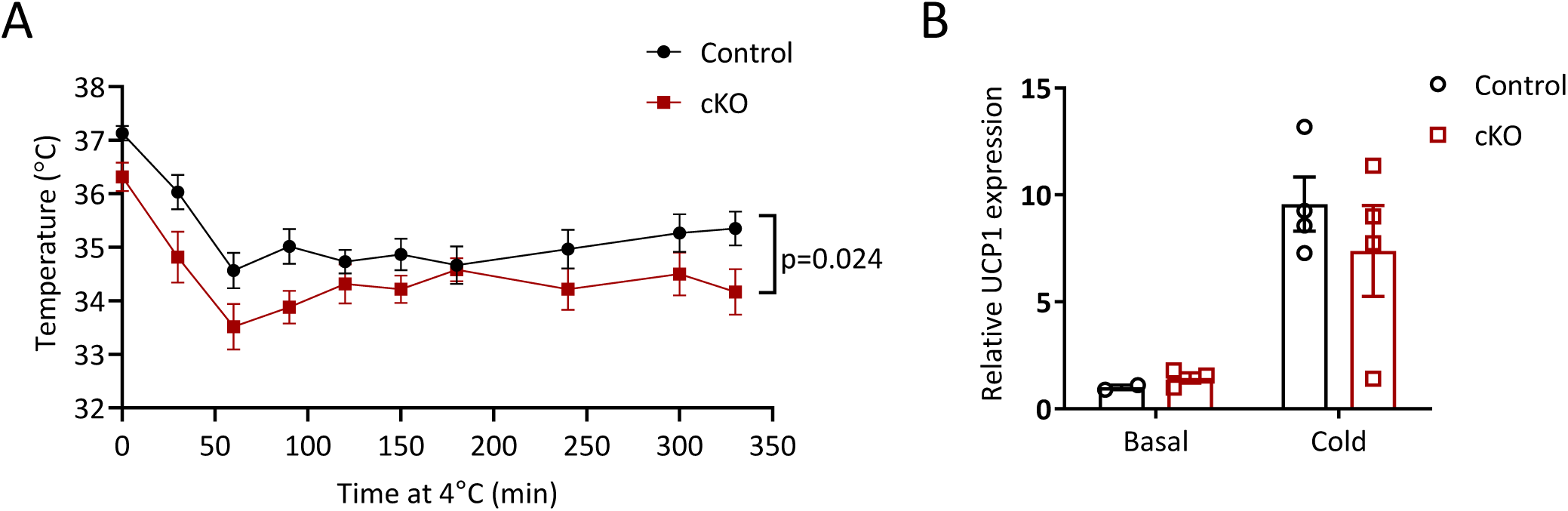
Otp loss in adulthood does not affect the mice’s ability to defend their body temperature under cold conditions. A. Subcutaneous temperature of control and Otp cKO mice under cold exposure (n=6). B. Relative UCP1 expression in the BAT of control and Otp cKO under basal conditions or following exposure to 4°C for 6h (n=2-4).

